# Stick of Sticks: Structural Features of the Amyloidogenic Peptide-DNA Complex

**DOI:** 10.1101/2024.11.05.622117

**Authors:** G.A. Arzamastsev, Y.A. Zabrodskaya, P. Garmay Yu., A.V. Shvetsov, N.S. Ivanova, D.S. Vinogradova, A.V. Arutyunyan, N.A. Verlov, V.S. Burdakov, T.N. Baymukhametov, A.L. Konevega, N.V. Gavrilova, O.I. Ivankov, E. Gorshkova Yu., V.V. Egorov

## Abstract

The search for peptides that can specifically bind to regulatory regions in DNA is a necessary step for creating drugs that can regulate gene expression. The study is dedicated to the peculiarities of binding of a model peptide, which carries an ionic self-complementary motif and can form amyloid-like fibrils [1], with model double-stranded DNAs. The stoichiometric ratios of the components of the complex were found using the retardation method in agarose gel. Using microscale thermophoresis, it was shown that the peptide in the amyloid-like state is capable of binding to model 45-bp double-stranded DNA, with a micromolar equilibrium dissociation constant. Using cryo-electron, transmission electron, and atomic force microscopy, the morphology of peptide-DNA complexes was studied. Using dynamic light scattering and nanoparticle tracking analysis, as well as small-angle neutron scattering, the spatial parameters of the resulting DNA-peptide complexes were characterized. Molecular dynamics simulations showed that the arginine side chains of the peptide are prone to interact with guanine nitrogenous bases. It was shown that the formation of peptide-dsDNA complexes interferes with the operation of restriction endonucleases that have guanine-cytosine pairs in the recognition center, which is consistent with the results of prediction of interaction sites obtained using computer modeling. The results of the work can be used in the development of peptides capable of interacting with functional regions of DNA, as well as in the development of new carriers for transfection of DNA constructs.

## 1. Introduction

The design of drugs based on substances capable of regulating gene expression is an urgent task [2]. A few structures of natural transcription regulators are known [3], [4]. Small synthetic peptides are also known that can specifically interact with certain DNA structural elements [5], [6]. Proteins and peptides capable of amyloid-like fibrillogenesis necessary to perform their functions are called functional amyloids [7], [8]. The prion-like properties of such polypeptides are also realized in processes where positive feedback is necessary - the reaction of a living system to small amounts of stimulus [9], [10]. Amyloid-like fibrils of a number of proteins are capable of interacting with DNA; DNA can also initiate conformational transitions necessary for the protein to acquire a beta conformation and oligomerization [11]. The formation of functional amyloids also occurs in some cases of epigenetic regulation. Recently, it was shown that similar mechanisms of expression regulation are realized both in yeast [12], and in mammals [13]. Of particular interest are supramolecular peptide complexes having a regular structure, especially, peptides capable of forming amyloid-like fibrils and of cooperative interaction with DNA [14]. Ionic self-complementary motifs are elements of the primary structure of polypeptides, including periodically located amino acid residues with alternating positive and negative charges of side chains [15], [16], [17], [18]. Although such motifs are often found in the functional domains of proteins, including DNA-binding ones, they are poorly represented in databases of proteins that modulate gene expression. Interestingly, in a review of the structural classification of DNA-binding proteins, only TATA-binding proteins have a beta conformation in DNA-binding motifs [19]. It should be noted that a number of such proteins are also prone to amyloid formation [20]. In addition, peptides bearing ionic self-complementary motifs have the potential to be used as therapeutic nucleic acid delivery vehicles. [21]. Previously we showed [18], that the GDIRIDIRIDIRG peptide is capable of forming amyloid-like fibrils. We also showed that GDIRIDIRIDIRG fibrils are capable of dissociation into oligomers in the presence of a guanine analogue, the antiviral drug Triazavirin [22]. Data obtained from molecular dynamics simulations in free diffusion mode indicated that the side chains of arginines are prone to interact with triazavirin heterocycles. We hypothesized that the interaction occurs through hydrogen bonding between the amines of arginine and Тriazavirin, as is the case with guanine [23]. Although guanine atoms in double-stranded DNA are involved in hydrogen bonding with cytosine, we hypothesized that perhaps the arginines in the peptide would be able to interact with guanine heterocycles.

The peptide GDIRIDIRIDIRG, which we previously studied, had low solubility, apparently due to electrical neutrality at physiological pH values. Therefore, despite the unique properties of such a peptide, the prospects for its use remained vague. In this work, we study a shortened version of the peptide GDIRIDIRIDIRG – IRIDIRIDIRG (called SID), which retains self-complementarity, but has a non-zero total charge at neutral pH and is more soluble. In this work, we study the structural prerequisites for the specific interaction of amyloid-like fibrils formed by a model peptide carrying an ionic self-complementary motif with model DNA sequences.

## 2. Materials and methods

### 2.1 Peptide IRIDIRIDIRG (SID)

Peptide with amino acid sequence IRIDIRIDIRG (called SID) 0was synthesized by “Verta” Research and Production Company (Russia), the purity is more than 95%. Reagents used were produced by Sigma Aldrich (USA) or New England Biolabs, Inc (USA). Double-stranded DNA (dsDNA) (1P, 45 base pairs) was synthesized by DNA Synthesis (Russia), and labeled with Cy5 along the plus strand at the 3’ end. The sequence of such DNA is given in Table 1. Lambda phage DNA was produced by SibEnzym (Russia).

**Table 1.**
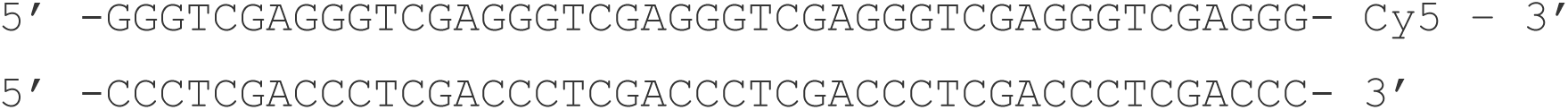
1P DNA sequence.

### 2.2 Preparation of fibrils from SID peptide

To obtain fibrils, the SID peptide was dissolved in 100% dimethylsulfoxide (DMSO; Dimexid, Tula Pharmaceutical Factory LLC, Russia), at a concentration of 10 mg/ml, then HEPES buffer (Sigma, USA) was added to a concentration of 1 mg/ml for the peptide; the mixture was incubated at 55°C for 180 min.

### 2.3 Preparation of peptide-DNA complexes

The peptide-DNA complexes under study were obtained by mixing dsDNA dissolved in HEPES buffer at the appropriate concentration with SID peptide dissolved in DMSO. The resulting mixtures were incubated at a temperature of 55 °C for 3 hours in a BioRad thermostat (USA).

### 2.4 Atomic force microscopy (AFM)

All reaction mixtures were diluted 10 times with a buffer for DNA binding (4 mM Tris-HCl (pH 7.5), 4 mM NiCl2) [24]. 10 μL of the diluted sample was applied to freshly cleaved 2SPI mica (USA) and incubated for 3 minutes at room temperature. The samples were then washed with deionized water and dried in a gentle stream of compressed air. AFM images were obtained in air using an atomic force microscope NT-MDT Solver BIO (Russia) with a Smena-B measuring head operating in semi-contact mode, using cantilevers NT-MDT NSG01, NSG03 (Russia), at a scanning frequency of 1 Hz and resolution 512x512 pixels. Surface topography of DNA and DNA-peptide complexes were analyzed using the NT-MDT software package. Images were processed in the Gwyddon software package [25].

### 2.5 Thioflavin T (ThT)

Amyloid-like fibrils were detected using ThT [26]. Thioflavin T (Sigma) was dissolved in phosphate buffered saline (PBS) and filtered through a 0.2 µm syringe filter. ThT concentration was then determined using an extinction coefficient of 36 mM^-1^ cm ^-1^ at a wavelength of 412 nm. To measure ThT fluorescence, samples of fibrillar complexes of the peptide and its complexes with lambda phage DNA were prepared in PBS buffer. 2 mL of 10 μM ThT solution was mixed with the peptide, model DNA, and DNA-peptide complexes and transferred to a cuvette for fluorescence measurement. ThT fluorescence was measured at room temperature using a Hitachi7000 fluorimeter with an excitation wavelength of 440 nm and recording at 450–700 nm.

### 2.6 Congo red (CR)

Congo red dye produced by Sigma Aldrich (USA) was used. A 500 μM (10-fold) dye solution in PBS was prepared and stored. Before measurement, a 5-fold molar deficiency of the sample (10 μM) was added to the dye diluted 10 times with PBS [25]. The absorption spectrum was obtained on a Hitachi 3310 instrument using disposable cuvettes with a volume of 100 μl.

### 2.7 Agarose gel retardation

Peptide fibrils at the appropriate concentration in HEPES buffer were added to double-stranded model DNA and incubated for 180 minutes at 55℃. Then samples were mixed with gel loading dye (New England Biolabs) and was applied to the 1.5% agarose gel wells (in TE buffer). Electrophoresis was carried out at 130 mA for 1.5 h at a constant temperature of 20℃. Visualization of the bends was carried out using the BioRad ChemiDoc gel documentation system (BioRad, USA) sequentially in the fluorescence detection mode of ethidium bromide and Cy5.

### 2.8 Microscale thermophoresis (MST)

To conduct an experiment to determine the binding constant of peptide monomers to DNA, samples of DNA complexes (1P) with peptide fibrils (SID) were prepared in various concentrations. DNA was dissolved in a 20 μl HEPES buffer solution to the final concentration of 50 μM. Next, a model peptide dissolved in 100% DMSO was added to each of the 16 samples in concentrations from 0.021 nM to 700 nM; the DMSO concentration in all samples was 2% v/v. To form complexes, the resulting mixture was left for 180 minutes at a temperature of 55°C. The equilibrium dissociation constant KD was calculated from the obtained curve (using equation [1])

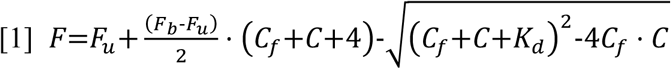

MST experiments were carried out at the National Research Center “Kurchatov Institute” – PNPI on a Monolith NT.115 device (NanoTemper Technologies GmbH, Germany) in accordance with the manufacturer’s protocols.

### 2.9 Dynamic light scattering (DLS)

Experiments to determine the hydrodynamic radius using DLS were carried out for DNA fragments (1P) 45 nucleotide pairs long in complex with the SID peptide. The formation of complexes occurred in standard HEPES buffer for 180 minutes at a temperature of 55°C. DLS measurements were carried out using a laser correlation spectrometer LKS03 (Intoks, Russia) equipped with a helium-neon laser with a wavelength of 633 nm. LKS03 software (Russia) was used for data collection and analysis. Measurements were carried out in disposable polystyrene cuvettes with an optical path length of 10 mm; the sample volume was 500 μl. Measurements were carried out at a distance of 4.65 mm from the cell wall with an automatic attenuator and at a controlled temperature of 25°C. For each sample, 5 measurements of 180 seconds were performed.

### 2.10 Nanoparticle tracking Analysis (NTA)

NTA measurements were performed on a NanoSight 2.3 (USA) equipped with a sample chamber with a 640 nm laser and a Viton fluoroelastomer O-ring. Samples were injected into the sample chamber using sterile syringes until the liquid reached the tip of the nozzle. All measurements were carried out at room temperature. NTA 2.0 Build 127 software was used for data acquisition and analysis. Samples were measured for 180 s in High Dynamic Range mode, which splits the captured video into two videos with independent shutter speed and gain settings. Scattered light modes (wavelength 640 nm) and excitation mode at a wavelength of 406 nm and registration of only fluorescent particles were used (using a buffer containing 10 μg/ml Thioflavin T). The average particle size and standard deviation values were obtained using NTA software.

### 2.11 Restriction analysis

Determination of the location of specific peptide binding sites on DNA was carried out using electrophoretic separation of DNA fragments obtained during enzymatic hydrolysis of DNA. To carry out the experiment, 3 enzymes were used: Ava1, Hind3, Nde1 in the corresponding rCutSmart and 2.1 buffers (New England Biolabs) at concentrations recommended by the manufacturer and lambda phage DNA. DNA fragments were visualized by fluorescent DNA staining using ethidium bromide after electrophoretic separation in a 1.5% agarose gel.

### 2.12 Small angle neutron scattering (SANS)

SANS spectra were obtained on the YuMO spectrometer located on the fourth channel of the IBR-2 reactor (Frank Laboratory of Neutron Physics, Joint Institute for Nuclear Research, Dubna, Russia). To measure scattering spectra using the SANS method, solutions of peptides and peptide–DNA complexes were prepared in D2O at a concentration of 1 mM. The measurements were carried out according to the standard method for this device, described earlier [27]. Scattering was recorded by two detectors simultaneously in the Q region from 0.006 to 0.3 Å.

Preliminary data processing was carried out according to the scheme described in [28]. Obtaining and normalizing the curves in absolute units was carried out using vanadium located directly in front of the detector. During the spectra analyses the following relationship should be fulfilled:

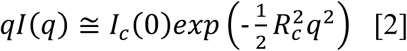

Based on this ratio, to determine the average radius of the resulting complexes, the spectra were plotted in Guinier coordinates for fibrillar structures (lnIq versus q^2^, Figure 11), after which a linear approximation of the initial sections of the scattering curves was performed using the least squares method (using the Origin2018 software package) and the slope angle for each of the lines was calculated.

From formula [2] it follows that the radius Rc can be calculated:

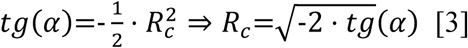

The Origin2018 software package was used to visualize the curves.

### 2.13 Molecular dynamics simulation

The results of computer modeling were obtained using the computing resources of the supercomputer center of Peter the Great St. Petersburg Polytechnic University (www.spbstu.ru).

As the initial state, we used fully unfolded conformations of the IRIDIRIDIRG peptide and unfolded 1P DNA in the B-form, generated in the PyMol program [29]. Using GROMACS [30] a system was built in the form of a cubic box with dimensions of 100^3^ Å^3^, containing 90 monomers of the SID peptide and Na^+^ and Cl^-^ atoms in a number corresponding to a 25 mM NaCl concentration. This ion concentration was used for modeling of the buffer ionic strength used in *in vitro* experiments. For peptides the field amber14sb was used [31], and tip3p for water [32]. The prepared starting configuration of the polypeptide was placed in a box with water molecules. The subsequent procedures were carried out similarly to those described in [33]

Minimizing the energy of the system was carried out using initial velocities calculated based on the Maxwell distribution (for 310K). The Berendsen thermostat and barostat model were used [34]. For solvent equilibration 5 ns simulation with immobile protein was performed. The Nose-Hoover thermostat [35], [36], [37], [38] and Parinello-Raman barostat [39], [40] were used for followed 10 ns equilibration. Molecular dynamics simulation was performed for 150 ns. The obtained data was processed using the numpy and matplotlib packages. Images of the simulated trajectory were processed in the PyMol software package [41].

### 2.14 Cryo-electron microscopy (cryo-EM)

For cryo-EM sample preparation a 3 μl of protein solution was applied to the Quantifoil R1.2/1.3 300 mesh Cu grid, which was glow-discharged for 30 s at 15 mA using PELCO easiGlow (Ted Pella, USA). The grid was plunge-frozen in liquid ethane using a Vitrobot Mark IV (Thermo Fisher Scientific, USA) with the following settings: chamber humidity 100%; chamber temperature 4 °C; blotting time 3 s; blotting force 0. Cryo-EM data were collected on a Titan Krios 60-300 transmission electron microscope (Thermo Fisher Scientific, USA) with a Falcon II direct electron detector (Thermo Fisher Scientific, USA) at the National Research Centre “Kurchatov Institute”. The microscope was operated at 300 kV with a nominal magnification of 75000x, corresponding to a calibrated pixel size of 0.86 Å, total exposure dose was 80 e−/Å2 and fractionated into 32 frames during 1.6 s exposure time. A total of 400 micrographs (Figure 1B) were collected and pre-processed using Warp software [42]. Filaments were picked using CrYOLO [43] and then inspected and corrected using RELION [44] GUI. To estimate helical parameters two sets of extracted amyloid segments were classified using 2D classification procedure in cryoSPARC [45]. The first set of ∼110,000 segments extracted with 512 px box size (0.86 Å/px) were used to estimate helical rise. The most detailed class-averages (Sup. fig. 1 A) demonstrates periodicity corresponds to distance between adjacent beta-strands of ∼4.7 Å, estimated using averaged power spectra (Sup. fig. 1 B). The second set of ∼60,000 segments extracted with 1600 px box size and downsampled to 400 px (3.44 Å/px) were used to estimate helical twist (angular distance between adjacent beta sheets). The class-average shows clear crossovers with a distance of about 580 Å (Sup. fig. 1 C) giving an approximate twist of -1.5 degrees. To generate an initial model [like described in[44]], some class averages from the first set of segments were selected (Sup. fig. 2 A,B). Further refinement of the initial model using ∼50,000 segments has not yielded a reliable high-resolution structure.

**Figure 1.**
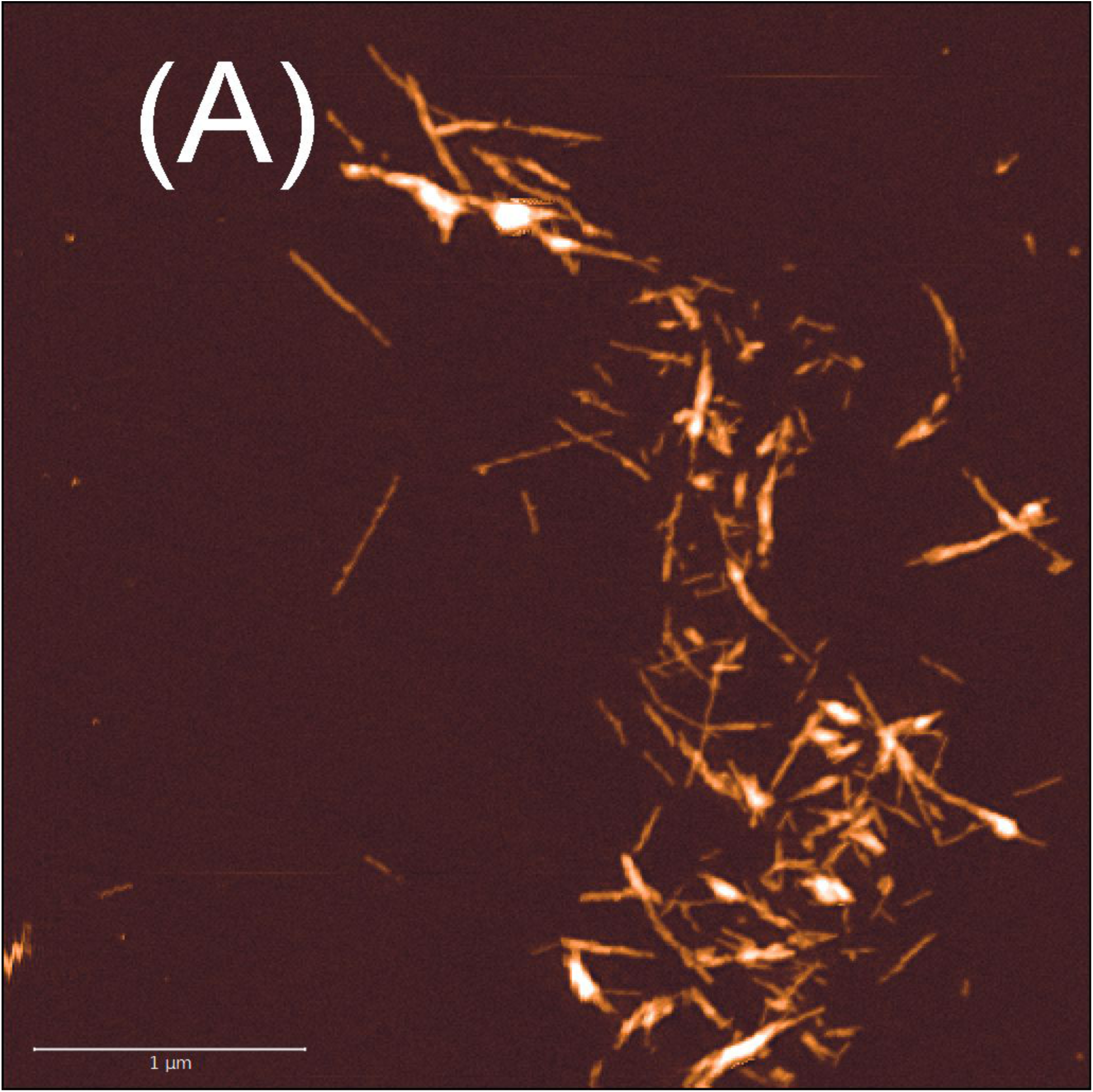
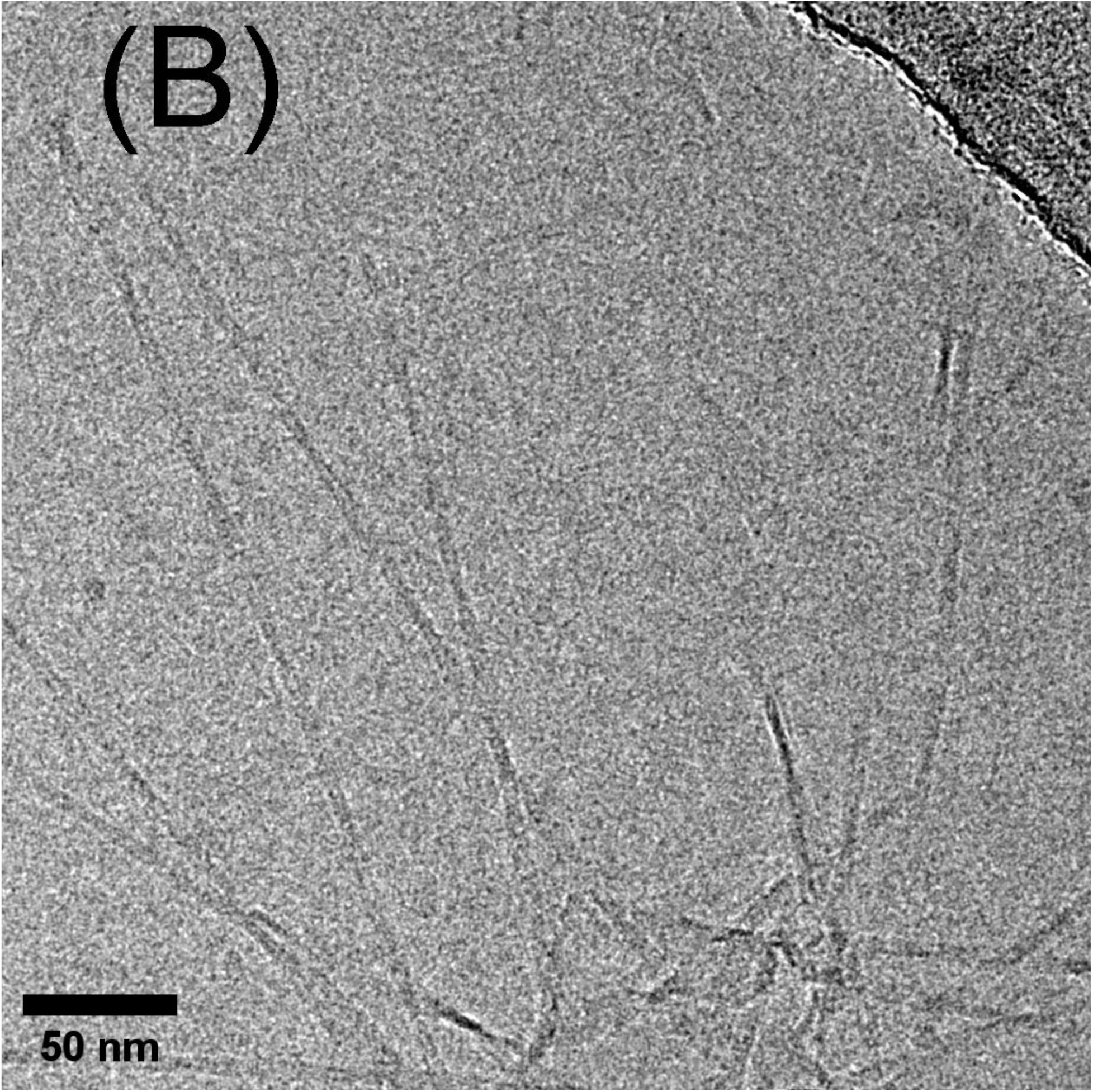
Microscopy of fibrillar structures formed by SID peptide. (A) Surface topography obtained using atomic force microscopy (scale bar is 1 um); (B) Representative row cryo-EM image of SID filaments (scale bar is 50 nm).

**Figure 2.**
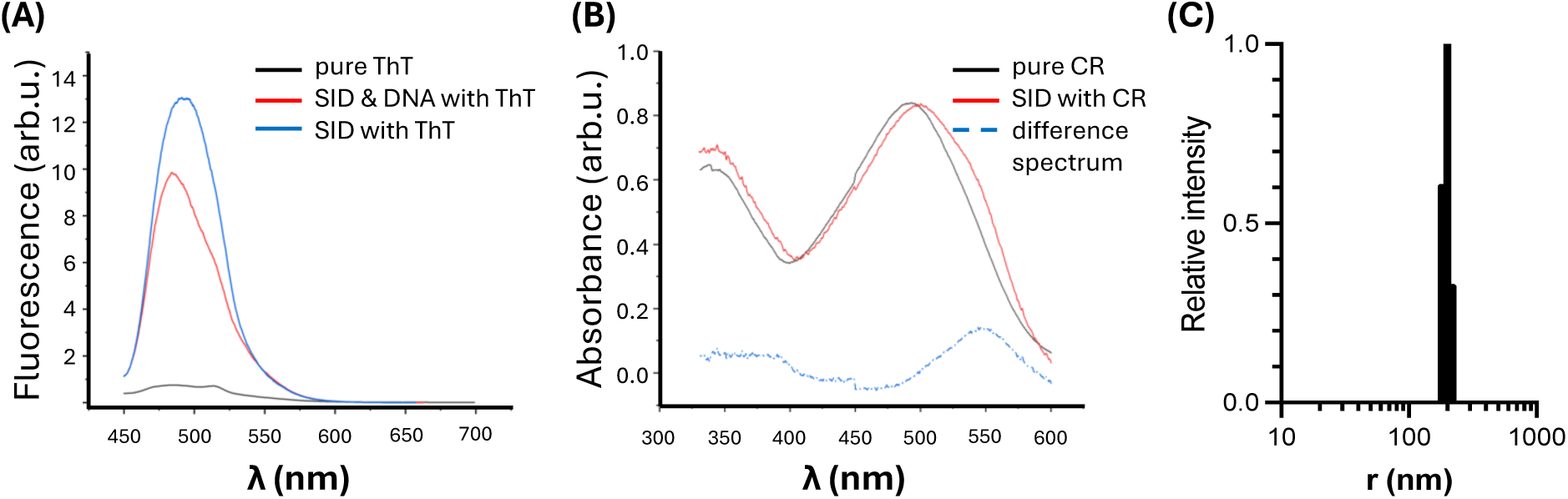
(A) - fluorescence spectrum of ThT when adding the model SID peptide to a solution; (B) absorption spectra of Congo red and its complexes with the SID peptide aggregates (C) Histogram of aggregate size distribution by dynamic light scattering in SID fibril solution prepared as described in 2.2 and diluted with HEPES to the concentration appropriate to measurement. The distribution of scattering intensity is given

### 2.15 Transmission Electron Microscopy (TEM)

For TEM, peptide or peptide-DNA complex solution samples were applied to carbon-coated copper grids. After adsorption of the proteins to the grid surface for 1 min, grids were washed twice with distilled water, then incubated with a 2% solution of sodium phosphotungstic acid salt for 1 min. After staining, the grids were dried and examined in a JEOL JEM 1011 transmission electron microscope at an accelerating voltage of 80 kV. Electron micrographs were obtained using a Morada digital camera (Olympus Inc.)

## 3. Results and discussion

### 3.1 Peptide SID (IRIDIRIDIRG) is capable of forming amyloid-like fibrils

To select SID peptide fibrillogenesis conditions, nanoparticle tracking analysis (NTA) in presence of thioflavin T (ThT) was performed (see Materials and Methods, sec. 2.10) using both registration of particle trajectories in transmitted light and only fluorescent particles upon excitation at 488 nm. Complexes obtained under the conditions described in Section 2.2 e were registered identically under both measurement conditions, indicating the presence in the solution of only particles capable of binding ThT, i.e., of an amyloid-like nature. In further experiments, only fibrils obtained under such conditions were used.

We initiated fibrillogenesis of the SID peptide as described in Material and Methods section. Figure 1 shows the results of atomic force (Fig. 1A) and cryo-electron (Fig. 1B) microscopy. In Figure 2 there are results of fluorimetry in the presence of ThT (Fig. 2A), spectrophotometry in the presence of Congo red (Fig. 2B). The morphology of peptide aggregates, the ability to form fluorescent complexes with ThT, as well as the ability of peptide aggregates to shift the absorption spectrum of Congo red to the right indicate that the truncated peptide IRIDIRIDIRG, just like the previously studied GDIRIDIRIDIRG, is capable of forming amyloid-like fibrils.

According to cryo-electron microscopy data (Figure S1, B) the distance between beta layers (estimated by Fourier classes with periodicity) is about 4.7Å, the distance between crossovers (helix pitch) is also estimated by direct measurement by one of the classes - about 580Å (Figure S1, C). Thus, the angular distance between two adjacent beta layers is about 3 degrees, which corresponds to the structures of amyloid-like fibrils. The fibril length measurements were made on the AFM image, using standard tools of the GWYDDION program on 20 free-lying fibrils shows that the average fibril length was 499±45 nm, structural data (thickness of fibrils) obtained from cryo-electron microscopy indicate that the fibril consists of maximum 2 twisted strands (Fig. S1, C). This structure is consistent with the published fibril structure of a similar peptide bearing an ionic self-complementary motif [46]. The typical distance between the peptide molecules in amyloid fibril is 4.7Å [47], that is, a protofibril with a length of around 500 nm represents about 2000 molecules stacked in parallel. We can estimate that one protofibril contains 24,000 monomer molecules (1338.81 Da). Together, these data allow us to estimate the molecular weight of one fibril as around 32000 kDa.

The calculated [48] hydrodynamic radius of fibrils (in the elongated particle model) was about 150 nm, thus the distribution of radii in the solution (Fig.2C) corresponds to the model obtained using cryo-electronmicroscopy.

### 3.2 Peptide SID is capable of forming complexes with short model DNA

The model DNAs used were 45 bp dsDNA (1P) labeled at the 3’ end with Cy5 (Table 1). The sequence of the model dsDNA 1P was chosen based on (i) the absence of self-complementarity, (ii) the high (80°C) melting temperature of double-stranded DNA, and (iii) the high content of GC pairs.

The presence of the binding was determined by DNA retardation in agarose gel. Using serial dilutions of peptide fibrils, the minimum concentration of the peptide at which no DNA entry into the gel was observed was determined (Figure 3). For 10 nM 1P DNA, this peptide concentration was 300 μM. Based on the results of the experiment, the ratio of the number of peptide molecules to the number of DNA molecules was calculated, which was 1 DNA molecule per 30,000 peptide molecules, which, according to structural data, approximately corresponds to 1 DNA molecule per 1 fibril.

**Figure 3.**
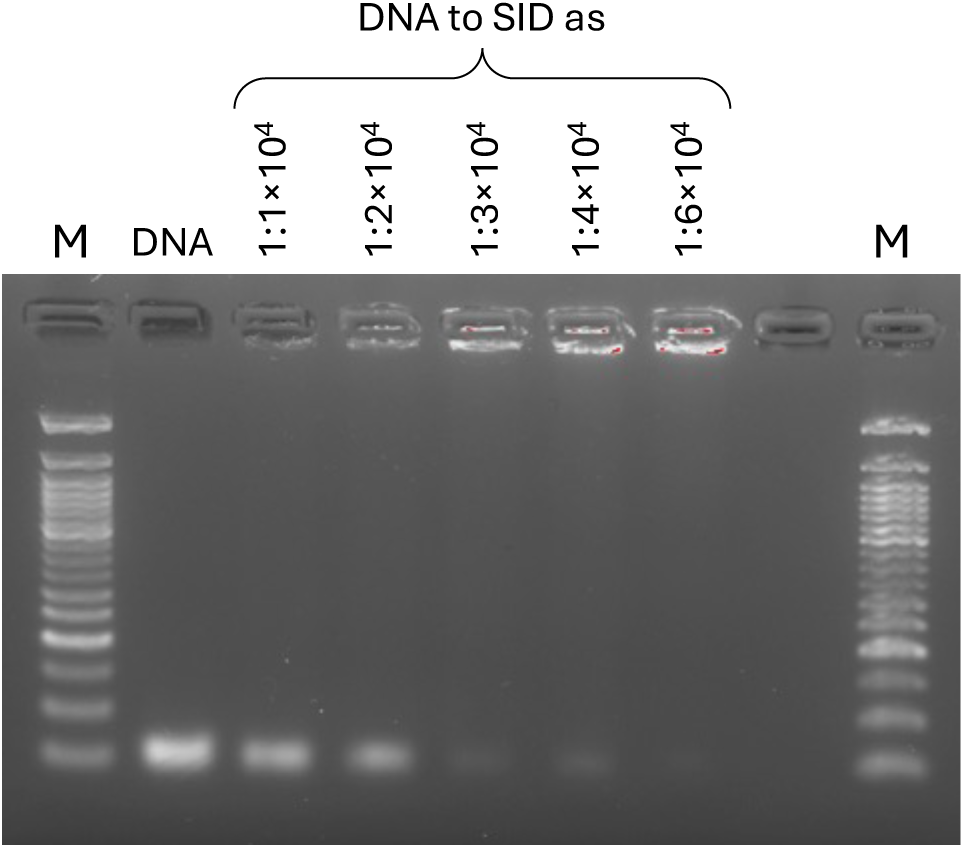
Titration of DNA 1P solution with SID peptide. M – 1000 kbase marker, «DNA» - DNA 1P 10nM, DNA:SID ratios used are designated above lanes

The study of the morphology of 1P DNA/peptide complexes by atomic force and electron microscopy, as well as by dynamic light scattering, did not show detectable differences between in morphology and hydrodynamic radii in such samples. The length of 45 bp DNA is about 17 nm (versus 500 nm of fibril length), and is shorter than DNA persistent length. 1P DNA is rigid and has no periodicity in structure which is characteristic for double-stranded DNA. We supposу that 1P DNA attaches to SID peptide fibrils.

It should be also noted that when using unlabeled DNA, 1P fibrils showed deterioration in binding to ThT (Figure 2A, red line), which may be a consequence of the coincidence of binding sites for DNA and dye on the fibril and competition between them. It is also possible that this is due to differences in the microstructure and ThT binding sites of the peptide fibrils formed in the presence and absence of DNA.

### 3.3 DNA-peptide equilibrium dissociation constant estimation

Microscale thermophoresis experiments were performed to determine the equilibrium dissociation constant and assess the specificity of peptide binding to model DNA. Sixteen samples were prepared, each containing a constant concentration of fluorescent DNA (50 μM) and concentrations of peptide ranging from 0.021 nM to 700 nM.

The data presented (Figure 4) show that when the peptide concentration reaches 5.4 nM, peptide-DNA complexes are formed. The calculated equilibrium dissociation constant KD was about 5±3*10^-8^ M, which corresponds to specific binding (10.1016/j.tibs.2020.04.005).

**Figure 4.**
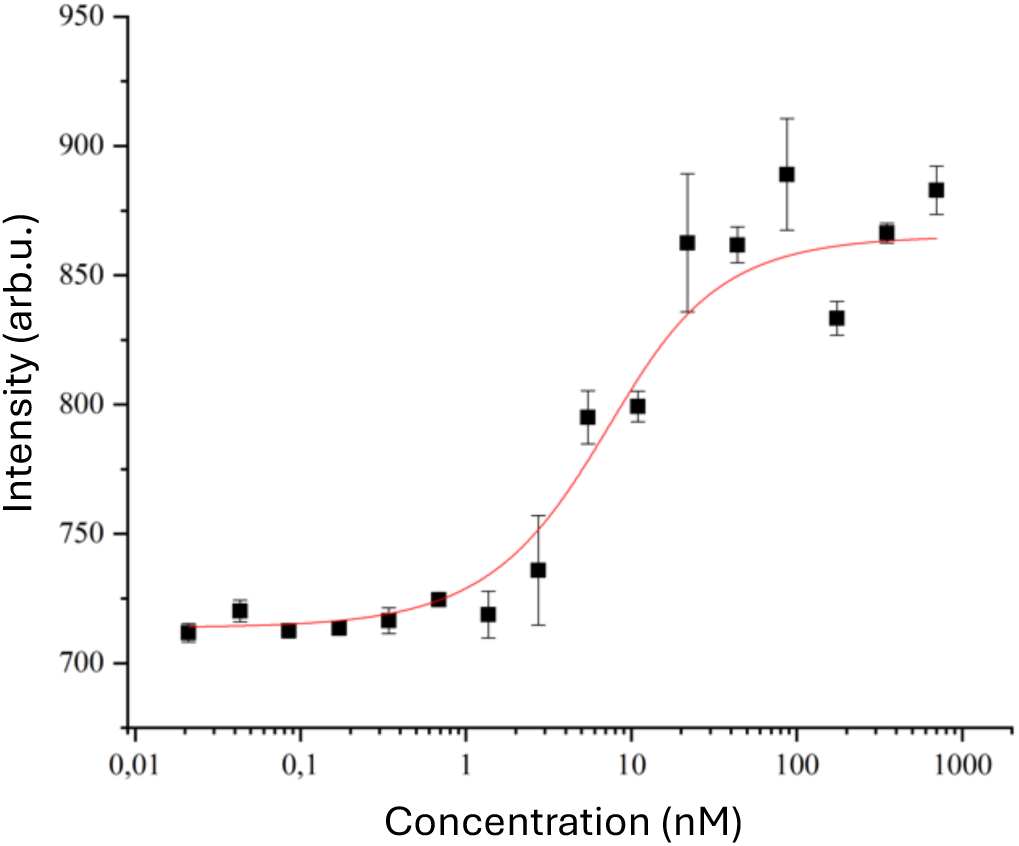
Plot of intensity versus concentration of SID peptide in the sample.

### 3.4 Mechanisms of interaction between the peptide and model DNA

To determine possible mechanisms of binding of the peptide to the model DNA, experiments on molecular dynamics simulation in the free diffusion mode were carried out. Figures 5 and 6 show the results of the analysis of the data obtained as a result of the simulation. As an example, Fig. 6 shows a heat map of the distances between nucleotides of the model DNA for two of the 90 peptides used in the simulation.

**Figure 5.**
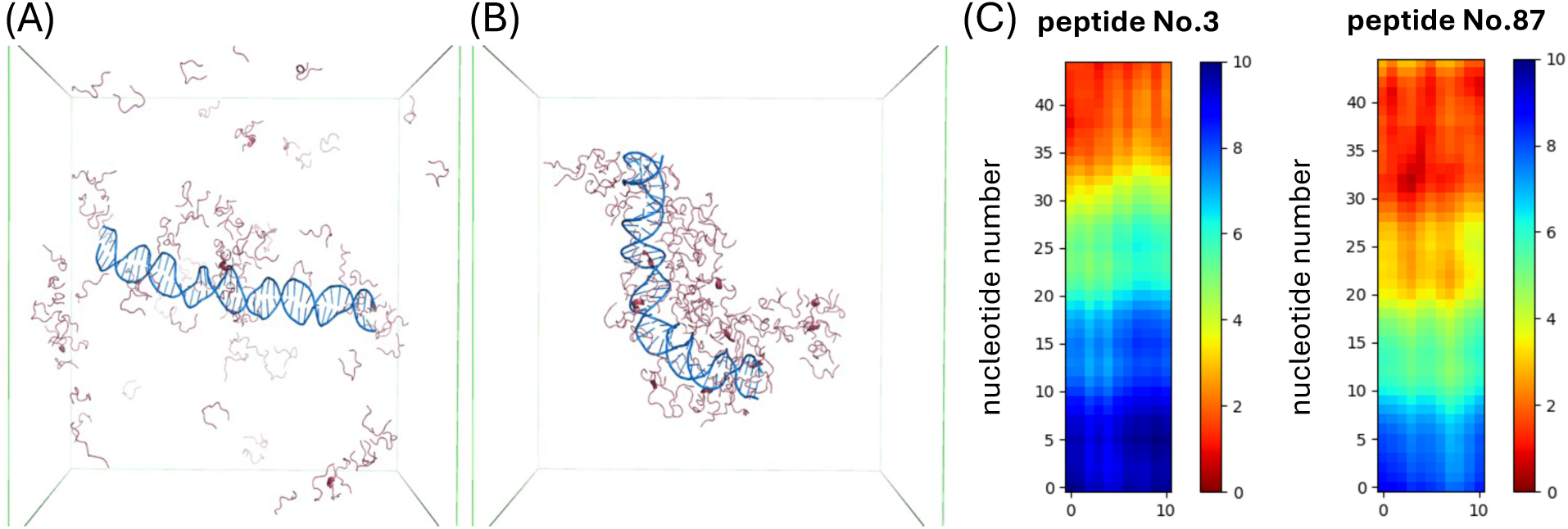
Sequential changes occurring to the system at 0 ns (A) and 150 ns (B) stages. The cube size is 10*10*10 nm. Panel C shows the example of heatmap of aminoacid residues to DNA distance distribution for two peptides from the simulation

**Figure 6.**
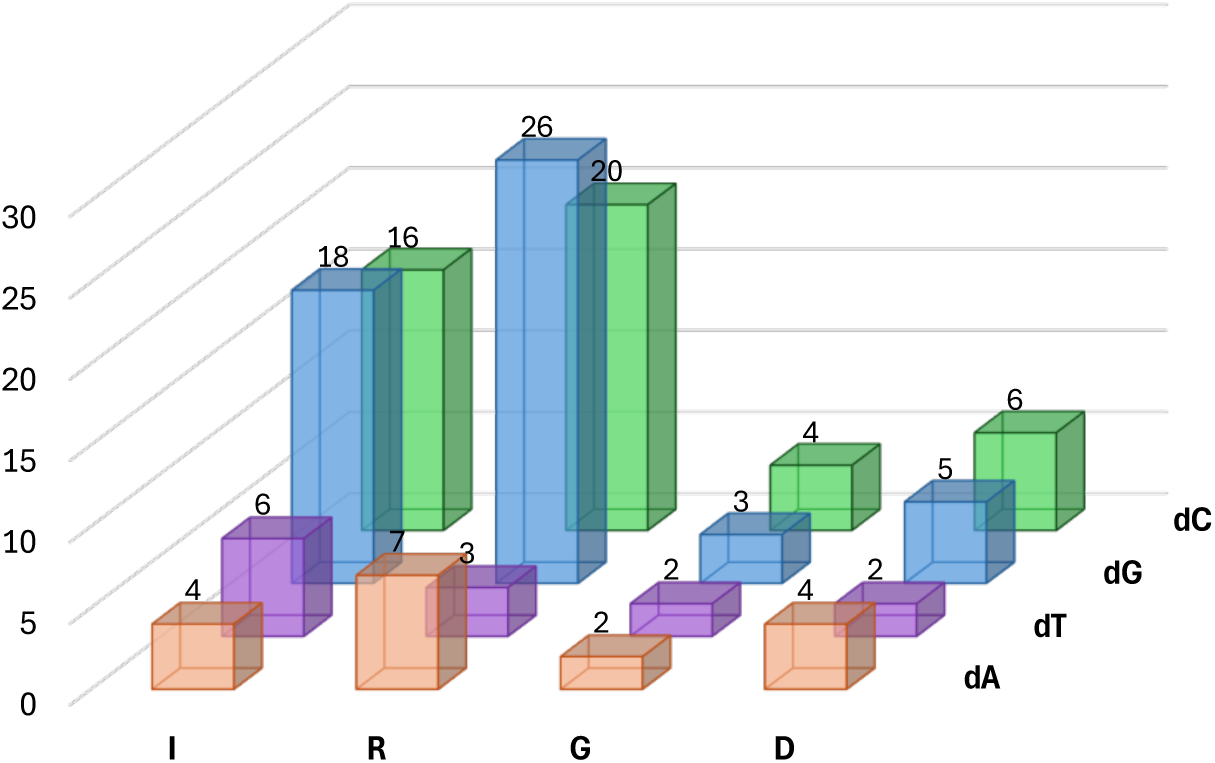
Maps of the modes of distances of amino acid residues of two randomly selected SID molecules No. 3(a) and No. 87(b) to the nitrogenous bases of model DNA 1P, during the simulation.

Based on the data presented in Table 2, a hypothesis was put forward about the selective interaction of the arginine and isoleucine residues of the SID peptide with GC-rich regions of DNA 1P.

**Table 2.** Number of contacts of nucleotides with amino acid residues of the peptide during the simulation process.

|  | I | R | G | D |
| --- | --- | --- | --- | --- |
| dA | 4 | 7 | 2 | 4 |
| dT | 6 | 3 | 2 | 2 |
| dG | 18 | 26 | 3 | 5 |
| dC | 16 | 20 | 4 | 6 |

The obtained data, despite the limitations of the model used (artificially close DNA and peptide molecules, the use of NaCl ions instead of Hepes buffer ions, the absence of DMSO), suggest that the side chains of arginines and isoleucines can enter into specific interactions with the nitrogenous bases of DNA.

### 3.5 Interaction of fibrils with phage lambda DNA

To test the hypothesis about the interaction of peptide molecules in the fibrils with GC-pairs of DNA, the lambda phage DNA was chosen as a model object. Conditions were selected for the formation of complexes of the SID peptide with the lambda phage DNA for subsequent restriction analysis using restriction enzymes that recognize GC- and AT-rich regions.

In experiments on retardation with lambda phage DNA (1 nM), the maximum concentration of the peptide that completely prevents DNA from entering the gel was 0.6 μM (Figure 7, 1:6*10^5^ ratio). This ratio was subsequently used to carry out the enzymatic hydrolysis reaction of DNA on the DNA-peptide complex.

**Figure 7.**
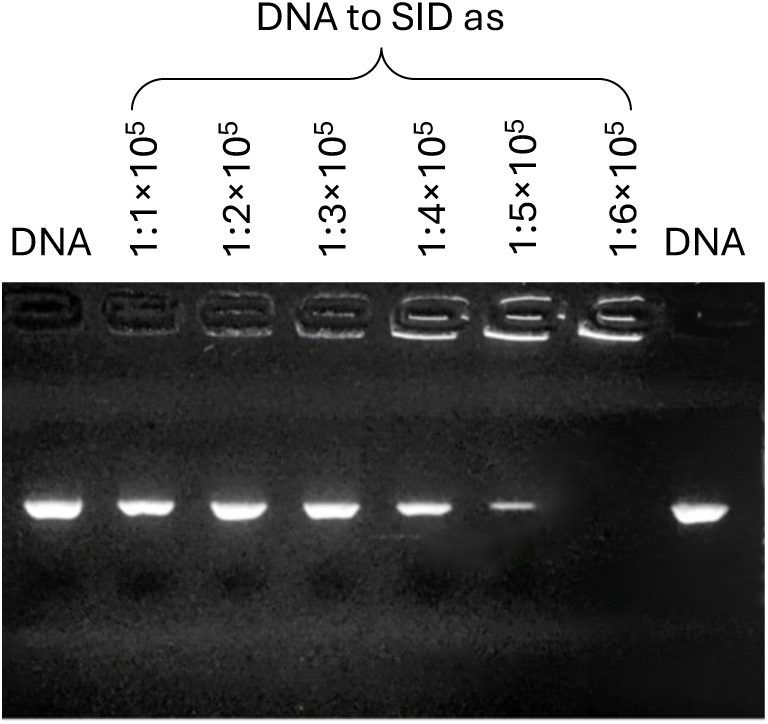
- Titration of lambda phage DNA solution with SID peptide. «DNA» -lambda phage DNA 1 nM, DNA:SID ratios used are designated above lanes

It should be noted that preliminary experiments to study the effect of concentrations of SID peptide comparable to concentrations of restriction endonucleases showed that it is not capable of directly inhibiting the enzymatic activity of endonucleases (data not shown).

To determine the binding sites of SID peptide to phage λ DNA, three different enzymes were selected: Ava1(CYCGRG), HindIII(AAGCTT) and Nde1(CATATG), catalyzing the reaction of DNA hydrolysis by specific sites noted in brackets.

The results of electrophoretic analysis of the products of the reaction of DNA hydrolysis of phage lambda are presented in Figure 8

**Figure 8.**
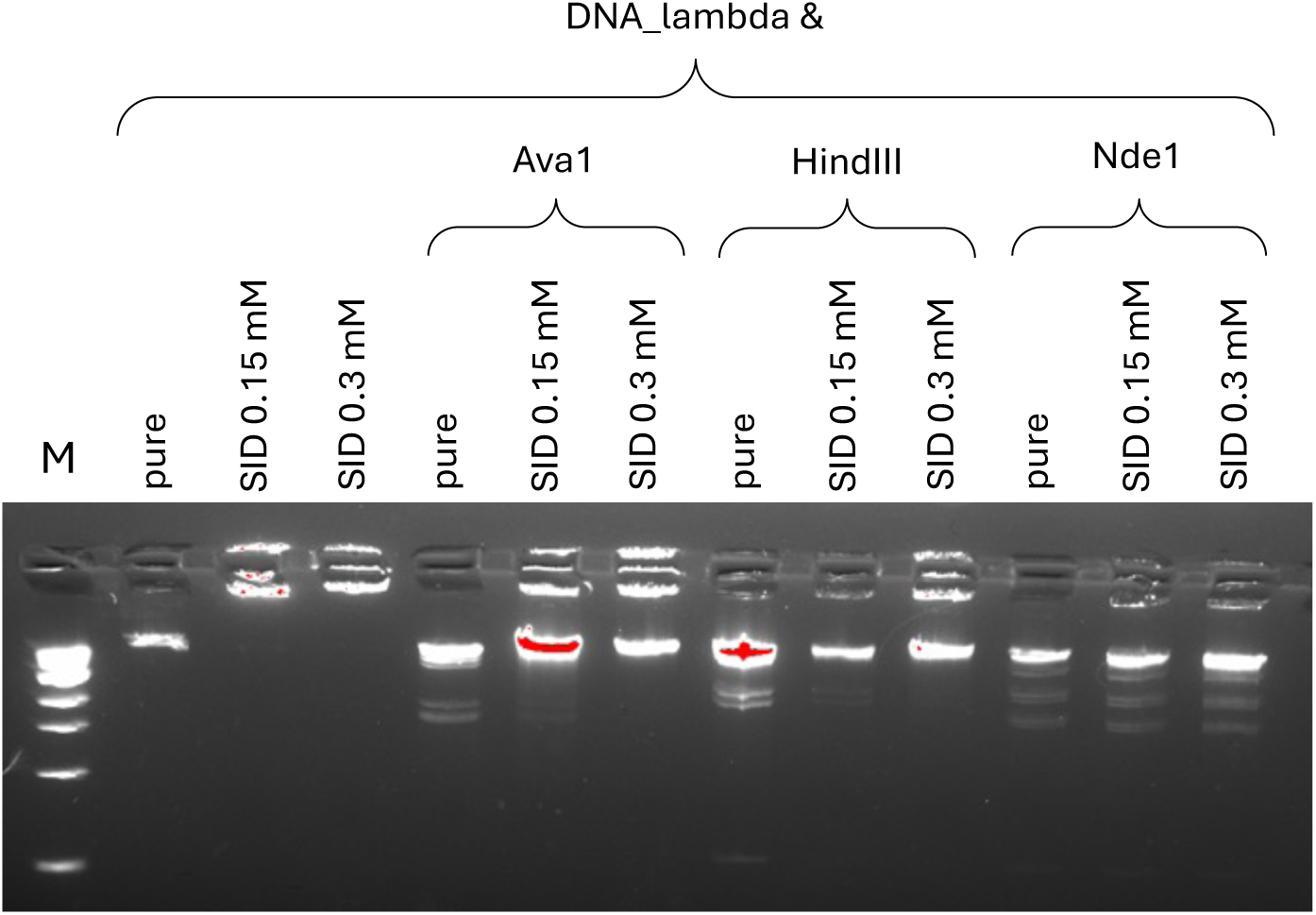
Result of electrophoretic separation of lambda phage DNA segments obtained using restriction enzymes Ava1, HindIII, Nde1 in the presence of SID. «M» – 1000kb marker, “pure” – phage λ DNA enzymes used and SID concentrations used are designated above lanes

The obtained results allow us to assume that inhibition of the hydrolysis reaction occurs only for enzymes that have specificity for recognizing GC-rich (Ava1(CYCGRG), HindIII(AAGCTT)) but not AT-rich regions (Nde1(CATATG)), which is consistent with the data obtained as a result of computer modeling and indicates the presence of selectivity in the binding of the peptide to the dsDNA molecule, depending on the nucleotide composition. Apparently, the peptide bound to DNA prevents the formation of the enzyme-substrate complex and protects the phosphodiester backbone of DNA from hydrolysis catalyzed by the enzyme.

#### Peculiarities of SID Peptide Binding to Short and Long DNA

For lambda phage DNA, the ratio preventing DNA from entering the gel is about 6*10^5^ peptide molecules per 1 DNA molecule, or about 20 fibrils per 1 DNA molecule, while for 1P DNA this ratio was 1 fibril per 1 DNA molecule. Comparison of the length of lambda phage DNA (about 16 μm) and SID fibril (about 0.5 μm) suggests that lateral interaction of fibrils and DNA may be involved in binding of lambda phage DNA, in contrast to binding to short DNA, where it seems that the attachment of short DNA to the fibrils is happening.. We hypothesized that different binding mechanisms may be involved in the interaction of DNA of different lengths with peptide, which are, however, based on the same mechanism of specific interaction with GC-rich regions. To test this hypothesis, an experiment was conducted to titrate already formed complexes of the

SID peptide/λ phage DNA with short 1P DNA followed by retardation in a gel. To conduct the experiment, the minimum concentration of the SID peptide was selected, leading to a complete delay of the λ phage DNA at the entrance to the gel, the concentrations of 1P DNA corresponded to those used earlier, during retardation of SID/1P complexes (Figure 9).

**Figure 9.**
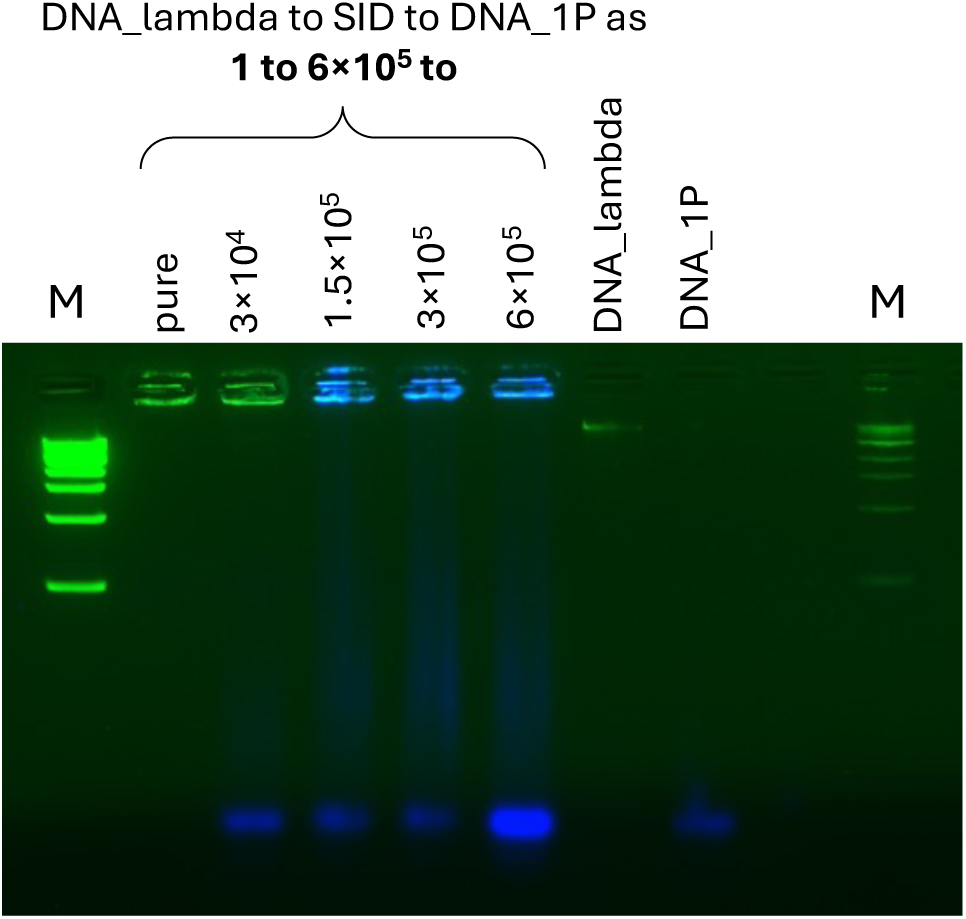
- Titration of SID – phage λ DNA complexes with 1P DNA solution. «M» - 1000base marker, «pure» - complex of lambda phage DNA with SID peptide 1/6*10^5^ SID concentrations used are designated above lanes

The results of the experiment showed that the capacity of peptide complexes for 1P DNA is less than 1:20 and does not significantly depend on whether it is in a complex with lambda phage DNA, i.e., the binding of long and short DNA by the SID peptide possibly occurs independently.

We performed negative-staining transmission electron microscopy of complexes of the SID peptide with double-stranded DNA of varying lengths. The microscopic images show that in the case of lambda phage DNA the peptide coats the DNA (Figure 10C), whereas with shorter DNA compact structures are formed, apparently consisting of individual peptide fibrils connected by DNA. In order to understand the structure of the complexes without being distorted by drying artifacts inherent in transmission electron microscopy, we studied the structure of the complexes in solution.

**Figure 10.**
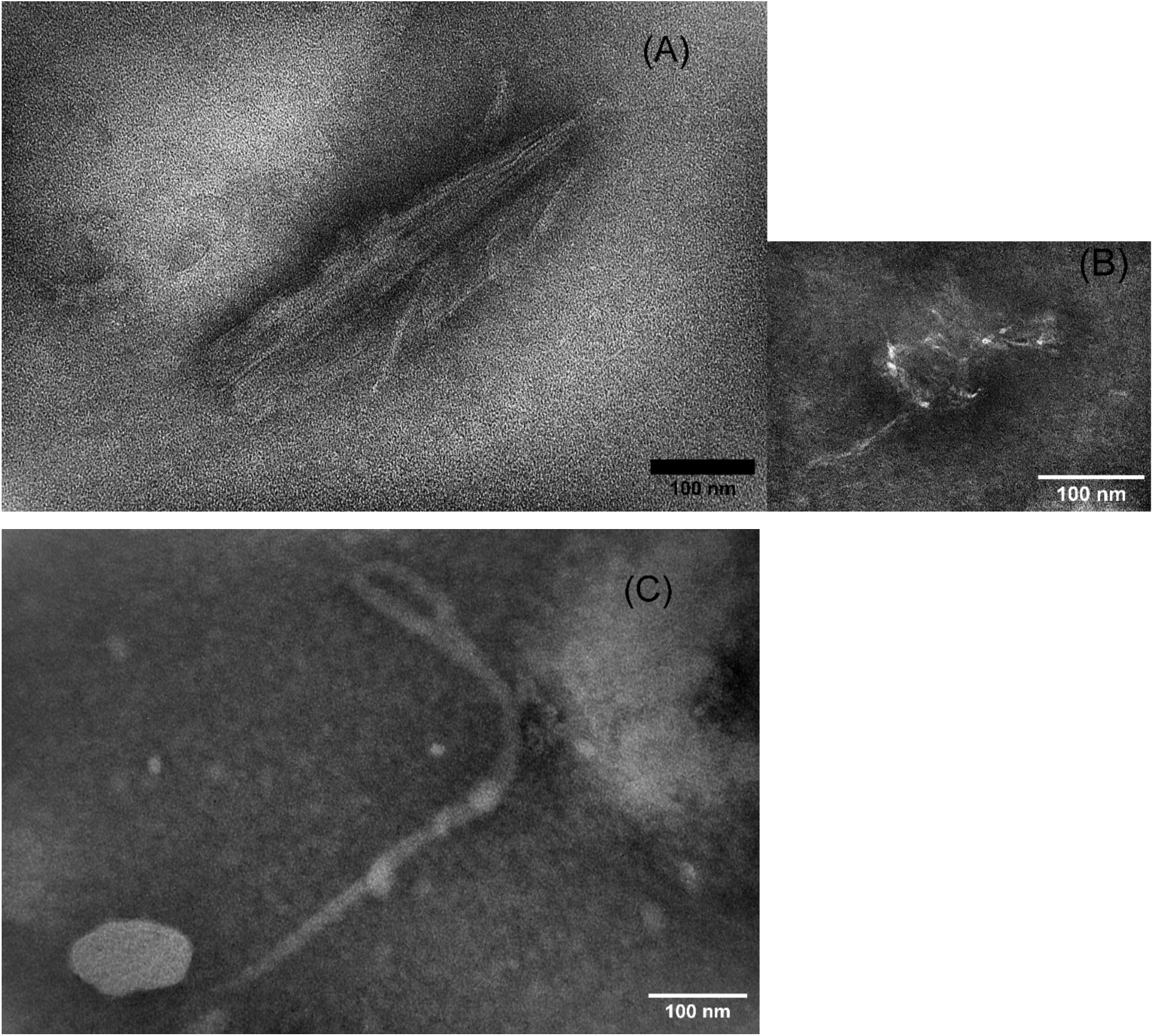
Electron microscopic images of SID peptide fibrils (A), SID peptide complexes with short double-stranded DNA (B) and with lambda phage DNA (C)

**Figure 11.**
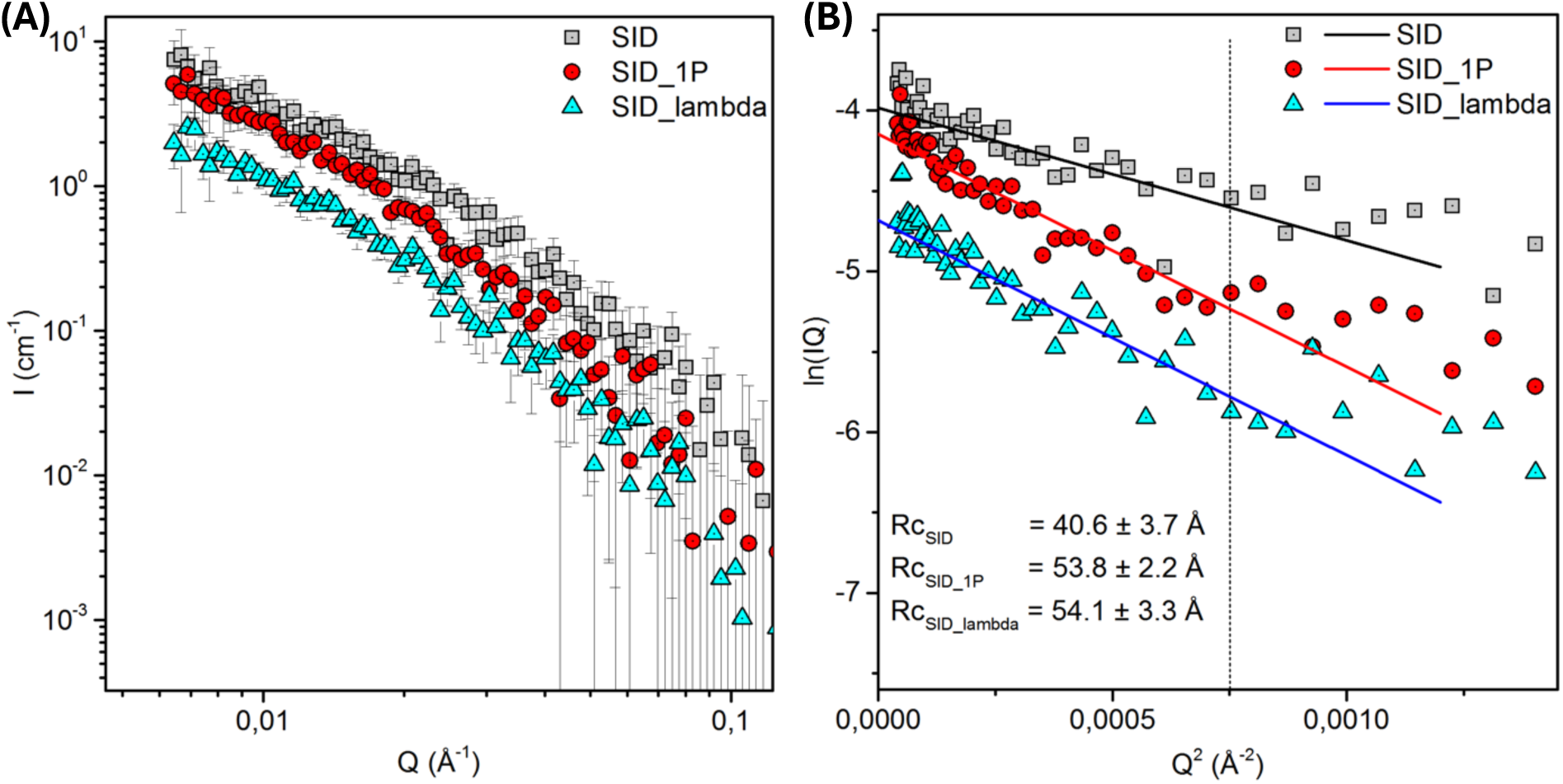
(A) Small-angle neutron scattering spectra in logarithmic coordinates. I is the scattering intensity, q is the scattering vector (B) SANS spectra on solutions of SID peptide, SID + 1P DNA, SID + phage lambda DNA, plotted in lnIq coordinates from q^2^ to determine the thickness of fibrillar structures. The dots show the experimental data, the lines show the linear approximation by the least squares method in the Guinier region (q*Rc < 1.3).

To compare structures of the DNA-fibrils complexes and peptide fibrils alone on small scales, small-angle neutron scattering (SANS) experiments were carried out on complexes of SID peptide fibrils with lambda phage DNA and 1P DNA. The obtained SANS spectra are presented in Figure 11.

Based on the nature of the spectra obtained, as well as on previous results obtained by the AFM method, it was concluded that in solution, the SID peptide itself and in combination with various types of dsDNA forms filamentous complexes of different radii.

Thus, the average radius of cylindrical structures formed in solutions of SID peptide and DNA of different lengths and nucleotide compositions was obtained (Figure 11 (B), Table 3). The obtained data satisfied the Guinier condition: q⋅R_C<1.3.

**Table 3.**
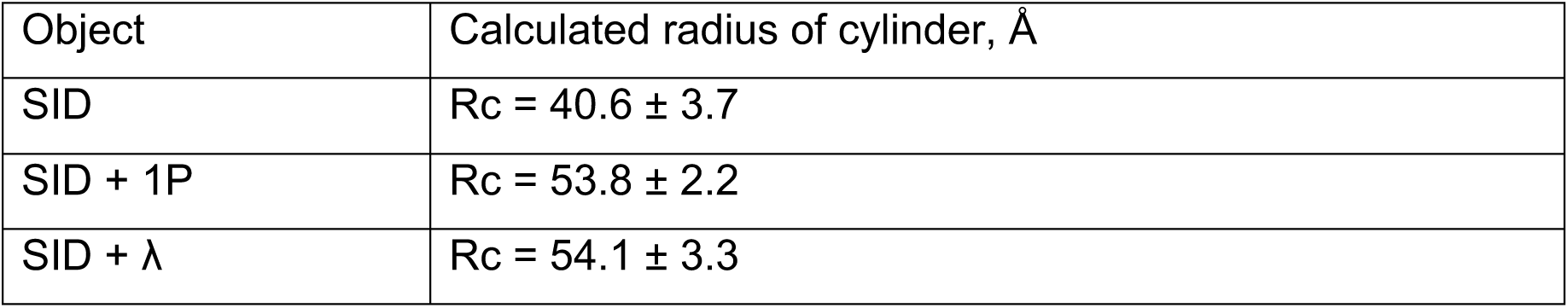
Calculated values of the cylindrical complexes radii.

| Object | Calculated radius of cylinder, Å |
| --- | --- |
| SID | $R_c = 40.6 \pm 3.7$ |
| SID + 1P | $R_c = 53.8 \pm 2.2$ |
| SID + $\lambda$ | $R_c = 54.1 \pm 3.3$ |

The average radius of the cylindrical complexes formed as a result of the interaction of the SID peptide with DNA 1P and phage λ differed from the average radius of the cylindrical complexes formed by the peptide itself by 10 Å, which is explained by the presence of DNA molecules in the complexes, which leads to an increase in the diameter of the cylindrical aggregates formed by the peptides by 20 Å.

The results of the experiment also showed that the structures of the complexes of lambda phage DNA with a peptide and 1P DNA with a peptide do not differ on scales of 6–60 nm, corresponding to the detectable range of SANS. It should be taken into account that the length of 1P DNA (17 nm) is less than the persistent length of DNA (45–70 nm, depending on the ionic strength), i.e. the difference in the binding mechanisms of 45 bp DNA and lambda phage DNA manifests itself on scales exceeding the persistent length and is associated with additional degrees of freedom arising due to the mobility of long DNA.

## Conclusions

We have shown that the IRIDIRIDIRG peptide carrying an ionic self-complementary motif is capable of forming amyloid-like fibrils. The peptide fibrils interact with a model 45 bp DNA, the equilibrium dissociation constant of such a complex is about 10^-7^ M. Although the mechanisms of interaction of the peptide fibrils with a short model DNA (45 bp) and a long model DNA (about 50 thousand nucleotide pairs) differ, the structural basis for the interaction of the peptide fibrils with DNA are GC pairs in DNA with arginine residues in the peptide. Thus, it appears that when interacting with long DNA, the fibrils coat it, while short DNA interacts with the surface of the fibrils. The resulting complexes can be used both for DNA delivery and as a basis for peptide drugs that are potentially capable of regulating gene expression.

## Authors’ contributions

Arzamastsev G.A. – manuscript writing, molecular dynamics, gel-electrophoresis; Zabrodskaya Y.A. — manuscript writing, SANS data analysis; Garmay Yu.P. — manuscript writing, atomic force microscopy; Shvetsov A.V. – manuscript writing, molecular dynamics; Ivanova N.S. – manuscript writing, atomic force microscopy; Vinogradova D.S. – manuscript writing, microscale thermophoresis; Arutyunyan A.V. – manuscript writing, dynamic light scattering; Verlov N.A. – manuscript writing, nanotracking experiments; Burdakov V.S. – manuscript writing, nanotracking experiments; Baymukhametov T.N. – manuscript writing, CryoEM experiments; Konevega A.L. – manuscript writing, microscale thermophoresis, Gavrilova N.V. – manuscript writing, TEM; Ivankov O.I. – manuscript writing, SANS experiments; Gorshkova Yu. E. – manuscript writing, SANS experiments, Egorov V.V. – manuscript writing, conceptualization, supervising

## Acknowledgements

The results of the work were obtained using computational resources of the supercomputer center in Peter the Great Saint-Petersburg Polytechnic University Supercomputing Center (www.spbstu.ru), SANS facilities of Joint Institute for Nuclear Research (project no. 2020-10-20-21-14-42), with financial support of National Research Center “Kurchatov Institute” (order No. 2557, October 28, 2021). MST experiments were supported by the Russian Science Foundation grant No. 22-14-00278 (to A.L.K.).

## Supplementary

**Fig. S1.**
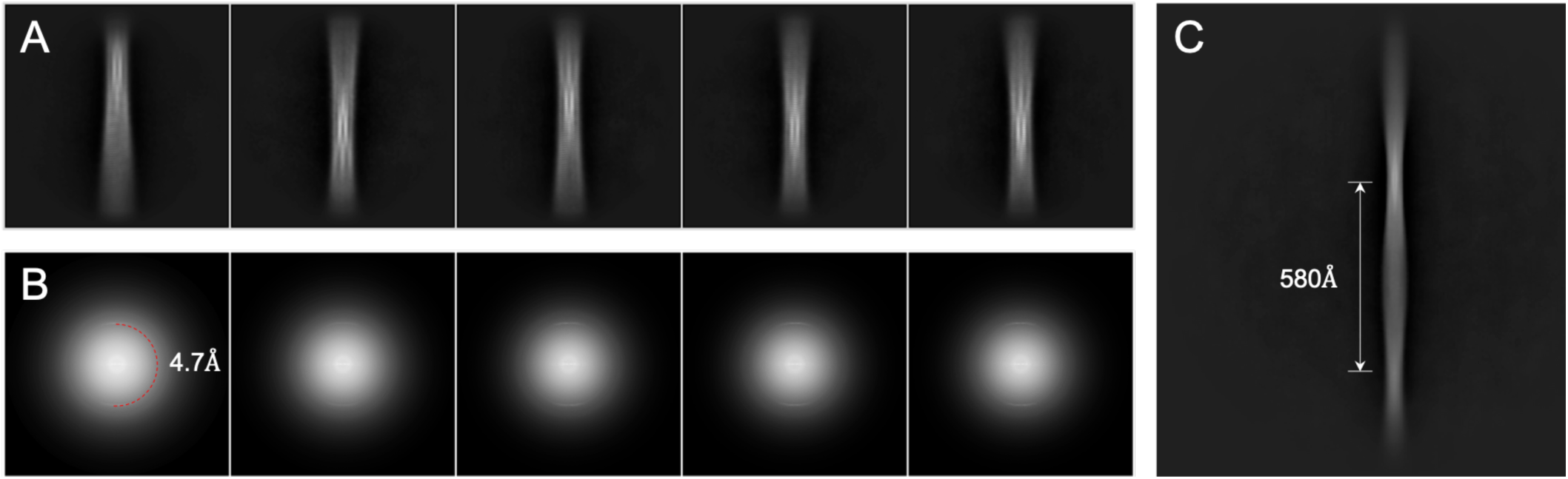
Representative 2D class-averages. A - Class-averages which clearly shows periodicity along the direction of the helical axis corresponding to the cross-beta quaternary structure. B - Correspondings averaged power spectra of segments in each class. C - Class-average which clearly shows the distance between crossovers.

**Fig. S2.**
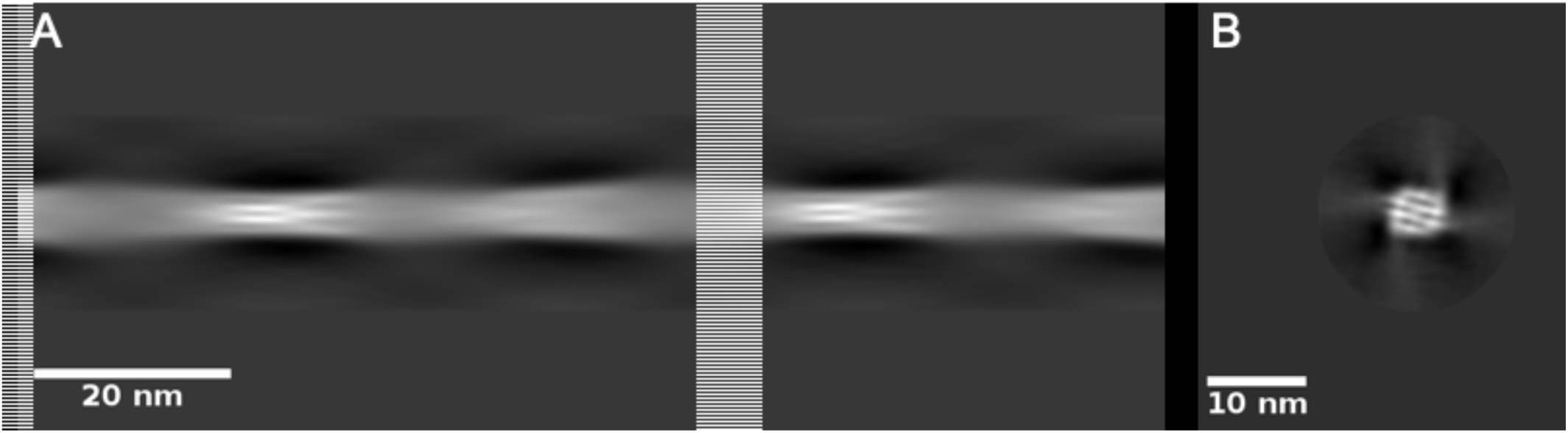
An initial model. A - View perpendicular to the helical axis. B - View along to the helical axis.

